# Antigen-driven CD8^+^ T cell clonal expansion is a prominent feature of MASH in humans and mice

**DOI:** 10.1101/2024.03.20.583964

**Authors:** Abbigayl E.C. Burtis, Destiny M.C. DeNicola, Megan E. Ferguson, Radleigh G. Santos, Clemencia Pinilla, Michael S. Kriss, David J. Orlicky, Beth A. Jirón Tamburini, Austin E. Gillen, Matthew A. Burchill

## Abstract

**Background and Aims:** Chronic liver disease due to metabolic dysfunction-associated steatohepatitis (MASH) is a rapidly increasing global epidemic. MASH progression is a consequence of the complex interplay between inflammatory insults and dysregulated hepatic immune responses. T lymphocytes have been shown to accumulate in the liver during MASH, but the cause and consequence of T cell accumulation in the liver remain unclear. Our study aimed to define the phenotype and T cell receptor diversity of T cells from human cirrhotic livers and an animal model of MASH to begin resolving their function in disease.

**Approach and Results:** In these studies, we evaluated differences in T cell phenotype in the context of liver disease we isolated liver resident T cell populations from individuals with cirrhosis and a murine model of MASH. Using both 5’ single cell sequencing and flow cytometry we defined the phenotype and T cell receptor repertoire of liver resident T cells during health and disease.

**Conclusions:** MASH-induced cirrhosis and diet-induced MASH in mice resulted in the accumulation of activated and clonally expanded T cells in the liver. The clonally expanded T cells in the liver expressed markers of chronic antigenic stimulation, including *PD1*, *TIGIT* and *TOX*. Overall, this study establishes for the first time that T cells undergo antigen-dependent clonal expansion and functional differentiation during the progression of MASH. These studies could lead to the identification of potential antigenic targets that drive T cell activation, clonal expansion, and recruitment to the liver during MASH.

## Introduction

The global rate of liver transplantation is rapidly increasing^1^, driven by an increased frequency of individuals presenting with liver disease due to poor diet and alcohol consumption^2–4^. A recent study demonstrated that approximately 14% of asymptomatic middle-aged individuals in the US had biopsy proven metabolic dysfunction-associated steatohepatitis (MASH)^5^. As such, MASH is a rapidly increasing epidemic that is predicted to affect 27 million people and cause approximately 78,000 deaths annually in the United States by 2030^6^.

Both innate and adaptive immune responses are critical for regulating steatosis and inflammation in the liver during the progression of chronic liver disease^7^. A key histological feature we^8^ and others^9^ have demonstrated in MASH and alcohol related liver disease (ALD) is a significant infiltration of T cells into the liver parenchyma. T cell accumulation can occur because of either antigen-independent T cell homing caused by increased chemokine production in the liver during MASH^10,11^, or antigen-driven T cell activation and subsequent clonal expansion. T cell clonal expansion in the liver has been reported in several diseases such as HCV^12,13^, AIH^14,15^, PBC^16,17^, and PSC^18^, but T cell clonal expansion has yet to be documented in MASH. However, disruption of T cell activation and effector function mitigates disease progression in different dietary models of MASH ^9,19^. While these studies demonstrated that CD8^+^ T cells promote liver injury and MASH, other studies have demonstrated a role for CD8^+^ T cells in promoting MASH resolution after diet reversal^20^ or in preventing MASH-induced hepatocellular carcinoma^21^. Therefore, T cell phenotype and function may not be static over the course of the disease and different T cell populations may have opposing roles in disease progression. As a result, the mechanisms by which T cell participate in MASH remains unresolved.

In this study we aimed to define the dynamics of T cell responses in MASH by probing the heterogeneity of liver infiltrating T cell subsets in both MASH-induced cirrhosis and in a mouse model of MASH. We demonstrate that both cirrhosis and a murine model of diet-induced MASH results in the accumulation of activated T cells in the liver. Specifically, we identified significant clonal expansion in the CD8^+^ T cell compartment in both humans and mice. The transcriptional profile of these clonally expanded T cells was associated with chronic antigenic stimulation. Together these data provide evidence that activated T cells clonally expand and accumulate in the liver via engagement of the T cell receptor in MASH.

## METHODS

### Patient Samples

Samples were selected from a biorepository of patients who had undergone liver transplantation and collected under the IRB protocol of MSK (Table 1). Non-diseased non-parenchymal cells (NPCs) were purchased from Triangle Research Laboratories (Lonza, Triangle research park, NC). All patients provided written and informed consent and the study was approved by the institutional review board at the University of Colorado—Anschutz.

**Table 1:** Patient Demographics.

| Tissue and blood samples |  |  |  |  |
| --- | --- | --- | --- | --- |
| <u>Sample</u> | <u>Age</u> | <u>Sex</u> | <u>Etiology of Cirrhosis</u> | <u>Race/Ethnicity</u> |
| ALD_1 | 58 | F | ALD | White |
| ALD_2 | 38 | F | ALD | White |
| ALD_3 | 61 | F | ALD | Hispanic |
| MASH_1 | 58 | M | MASH | White |
| MASH_2a/b | 49 | F | MASH | White |
| ND_1 | 97 | M | N/A | White |
| ND_2 | 35 | F | N/A | White |
| PBMC_1 | 35 | F | N/A | White |
| 10x genomics dataset samples |  |  |  |  |
| <u>Sample</u> | <u>Source</u> | <u>Dataset name</u> |  |  |
| PBMC_2 | 10x Genomics | PBMCs of a Healthy Donor (Next GEM v1.1) |  |  |
| PBMC_3 | 10x Genomics | Human PBMC from a Healthy Donor, 10k cells (v2) |  |  |
| PBMC_4 | 10x Genomics | 10k Human PBMCs, 5' v2.0, Chromium Controller |  |  |

### Animal Studies and Feeding Regimens

All mice used for experiments were 6 to 8-week-old male *C57BL/6* mice purchased from Charles River Laboratories (Wilmington, MA) or Jackson Laboratories (Bar Harbor, ME). Mice were randomly allocated to either a chow control or HFHC diet (#D09100301) made by Research Diets Inc. (New Brunswick, NJ). Mice were provided with indicated diet ad libitum for 25-31 weeks. All animal studies were approved by the University of Colorado Anschutz Institutional Animal Care and Use Committee.

### Liver digestion, NPC isolation and T cell enrichment

#### Human

Portions of diseased livers were harvested after transplantation and NPCs were isolated and frozen in a single cell suspension as described^8^. To enrich T cells from hepatic NPCs or viably frozen peripheral blood mononuclear cells (PBMCs)^22^, frozen samples were thawed in RPMI 1640 (Corning, NY,) containing 10% Human Serum AB (Gemini Bio-products, West Sacramento, CA) and 1% DNASE (MP Biomedicals, Santa Ana, CA). Cells were washed 2x with PBS containing 2% FBS (Atlas Biologicals, Fort Collins, CO). Viable T cells were enriched using the Easy Sep T cell isolation Kit followed by the Easy Sep Dead Cell removal Kit (Stem Cell Technologies, Vancouver, BC).

#### Mouse

Livers were digested, and NPCs were isolated as previously described^23^. Briefly, livers were perfused with 10mL HBSS (Gibco, Waltham, MA) prior to extraction and digestion with Collagenase Type IV (Worthington-Biochemical, Lakewood, NJ). NPCs were isolated using a 40% OptiPrep density gradient (Sigma-Aldrich, Burlington, MA), filtered, and washed. Following liver isolation and digestion, viable T cells were enriched using the MojoSort Mouse CD3 T Cell Isolation Kit and Mouse Dead Cell Removal Kit (BioLegend, San Diego, CA).

### Single cell (sc)RNA-sequencing preparation and analysis

Approximately 20,000 enriched T cells per sample were loaded on a 10x Genomics chromium controller, cDNA libraries were split, and samples were prepared using the Chromium Next GEM Single Cell 5’ Kit V2 or the Chromium Single Cell Human/Mouse TCR Amplification Kit (10x Genomics, San Francisco, CA) per manufacturer’s instructions. cDNA libraries were sequenced using a NovaSeq6000 (Illumina, San Diego, CA) to a depth of 50,000 reads for RNA and 4,000 reads for T cell receptors (TCRs) at the University of Colorado Genomics Shared Resource Core. 10x Genomics Cell Ranger was used for scRNA-seq upstream preparation as previously described^8^. Further analysis was performed using R (http://www.r-project.org). The R package DeContX^24^ was used to remove ambient RNA contamination and DoubleFinder^25^ to remove doublets. Cells with <500 detectable genes, >10% UMIs derived from mitochondrial genes for human T cells and >5% for mouse T cells, and clusters containing no *Cd3e* expression were removed from further analysis. TCR sequencing data was analyzed with scRepertoire^26^ and combined with scRNA-seq data. Seurat^27^ was used for all other downstream analysis including batch correction, with the exceptions of scTriangulate^28^ for unbiased cluster assignments and Presto^29^ for differential gene expression testing. For human T cells, DecoupleR^30^, using the multivariate linear model, was utilized to estimate transcription factor activity of CollecTRI^31^ transcription factor regulatory networks, and grouping of lymphocyte interactions by paratope hotspots (GLIPH)^32^ was preformed to cluster T cells predicted to bind the same peptide/MHC complex. For mouse T cells, fast Gene Set Enrichment Analysis (fGSEA)^33^ was used for gene set enrichment analysis and additional Ingenuity pathway analysis (IPA) was performed using IPA software (Qiagen, Germantown, MD).

### Quantification of Liver Pathology

For quantification of liver histology, a lobe of liver was fixed in neutral buffered formalin for 24 hours, embedded in paraffin wax and sectioned on glass slides. Liver sections were stained with hematoxylin and eosin (H&E) and scored semi-quantitatively for steatosis and inflammation as previously described^23,34,35^. For analysis of fibrosis, samples were stained with picrosirius red (PSR), and the total pixels/sample was quantitated using polarized light. Histologic scoring of liver sections was performed by D.J.O., who was blinded to the identity of the liver section being scored.

### Flow cytometry

Following liver isolation and digestion, samples were washed with FACS buffer and stained with surface antibodies (Supplemental Table 1) for 30 minutes at 4°C. Intracellular staining was performed using the eBioscience FoxP3/Transcription Factor Staining Buffer Set and protocol (Invitrogen, Waltham, MA). Stained cells were run on a CytoFLEX LX (Beckman Coulter, Brea, CA) following calibration of compensation with single stain samples. Results were processed with FlowJo software (Tree Star, Ashland, OR).

### Statistical analysis

Sequencing data statistical analysis of differentially expressed genes for fGSEA and IPA was preformed using the R package Presto^29^. IPA gene inputs were filtered to only include an adjusted P value of ≤ .01. Statistical analysis of flow cytometry data was performed in GraphPad Prism 10 (Boston, MA). A student’s t-test was performed for comparisons between two groups and 1-way ANOVA for comparisons between more than two groups. P-values were categorized as follows: *P<=0.05, **P<= 0.01, ***P<=0.001, ****P<=0.0001.

## Results

### Single cell mRNA sequencing identifies liver resident T cell populations in homeostasis and cirrhosis

A key histological feature of cirrhosis, regardless of etiology, is the accumulation of immune cells, particularly T lymphocytes^8^. However, in diseases such as MASH and ALD it is unknown whether these T cells are accumulating in the liver due to antigen-based activation and clonal expansion, or if they home to the liver independent of antigenic stimulation. To answer this question, we performed scRNA-seq on enriched T cells from the livers of individuals with cirrhosis resulting from MASH (n=2) or ALD (n=3) and from individuals with no noted liver disease (ND, n=2). As a control for liver dependent effects, we analyzed T cells from the peripheral blood of healthy control individuals (PBMC, n=4) and as a control for batch-dependent effects, we interrogated T cells from the same individual twice (MASH_2a, MASH_2b). Following T cell enrichment, cells were subjected to droplet-based 5’-end 10x Genomics scRNA-seq and sequenced cells were then filtered for *CD3E* expression (Supp. Figure 1A-C). Using unsupervised clustering and Uniform Manifold Approximation and Projection (UMAP), we visualized 25 unique clusters of T cells in the liver and blood that were represented in each individual sample and contained a unique transcriptional profile (Figure 1A,B, Supp. Figure 1D,E). Utilizing known markers of T cell functional state, we were able to identify multiple subsets of T cell populations including two clusters of Naïve/central memory (Tcm) (*SELL, CCR7, CD27, IL7R, LTB,* and *LEF1*), four clusters of resident memory (Trm) (*IL7R* and *CD69)*, twelve clusters of effector/effector memory (Tem) (*NKG7, GZMA, GZMB, CCL4,* and *XCL1)*, three clusters of gamma delta (γδ) T cells (*TCRD)*, and single clusters of T helper 17 (Th17) (*TIMP1, RORC*), T-regulatory cells (Treg) (*FOXP3, IL2RA, CTLA4*), TEM re-expressing CD45RA (Temra) (*NKG7, GZMK, CX3CR1*), and mucosal-associated invariant T cells (MAIT) (*ZBTB16, SLC4A10, TRAV1-2*) (Figure 1C,D)^36,37^. Unsurprisingly, liver resident T cells had a significantly different transcriptional profile from T cells isolated from peripheral blood, with liver resident T cells being almost exclusively contained in the CD8^+^ Trm, Tem, and Temra clusters and PBMC T cells being highly enriched for Naïve/Tcmand CD4^+^ clusters (Figure 1E). Surprisingly, we did not observe significant changes in the frequency of T cells in each cluster in the liver resident T cells between either ND and cirrhosis or between ALD and MASH (Figure 1E and Supplemental Figure 2). However, when investigating the transcriptional similarities between samples via hierarchical clustering of variable genes, we found that T cells from the majority of diseased livers (4 of 5) were much more similar to each other than to T cells from ND livers and PBMC derived T cells formed their own unique cluster (Figure 1F). Furthermore, analysis of predicted transcription factor activity using DecoupleR inference^30^ of collecTRI regulatory networks^31^ identified several regulons associated with T cell activation in cirrhosis. T cells from the peripheral blood had predicted transcription factor activity associated with naïve or effector T cells (FOXO1, MNT, and LEF1), whereas liver resident T cells from individuals without disease had transcription factor activity associated with effector or memory T cells (ID3, EOMES, and TCF7), and cirrhotic liver resident T cells had transcription factor activity associated with T cell dysfunction (ETV6, NFATC3, RBPJ, and HPB1)^38^(Figure 1G). These data demonstrate that cirrhosis results in the accumulation of activated and transcriptionally differentiated T cells in the liver.

**Figure 1:**
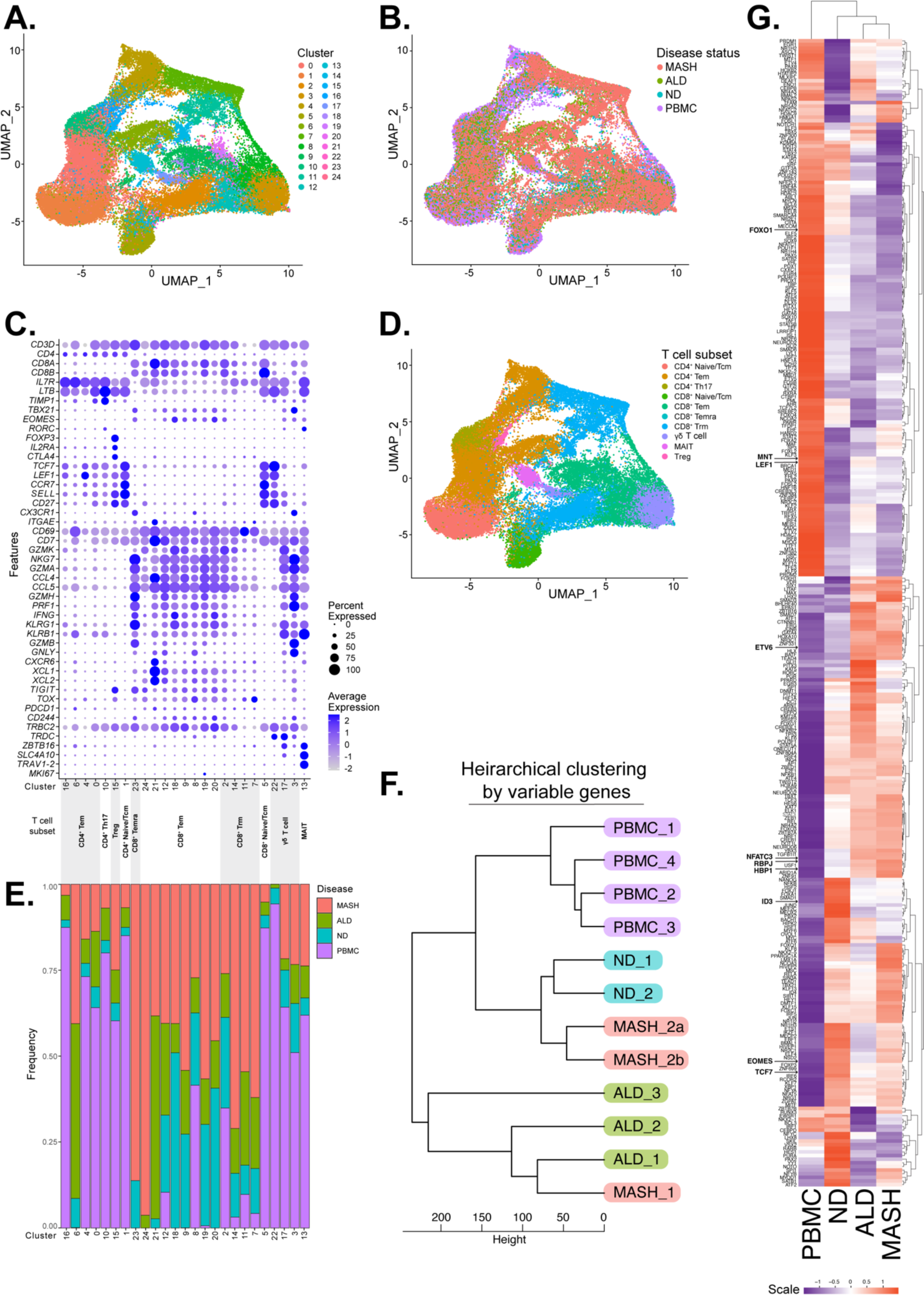
scRNA-sequencing of human T cells reveal transcriptional and inferred proteomic differences between healthy and cirrhotic individuals. (A) UMAP plot of 51,624 human T cells colored by scTriangulate clusters. (B) UMAP plot colored by disease status of T cell sample origin. (C) (top) Dot plot of selected gene (Features) expression grouped by cluster assignment. Larger dot size represents larger percent of cells expressing each feature and darker dot color represents the average expression level of cells expressing each feature. (bottom) T cell subsets as determined by select feature expression by cluster assignment. (D) UMAP plot colored by T cell subset assignment. (E) Bar graph displaying the proportion of cells within each cluster from each disease status. (F) Dendrogram of Euclidean distances between each sample as calculated from the expression levels of the top 2000 variable genes. (G) Heatmap of inferred transcription factor activity using DecoupleR inference of CollecTRI regulatory networks. Red represents higher inferred transcription factor activity while purple represented lower inferred activity. Top and Right dendrograms represent similarities between samples based on inferred activity or transcription factors, respectively.

### MASH-induced cirrhosis results in the recruitment of clonally expanded, chronically stimulated T cells to the liver

The most accurate way to determine if T cell accumulation results from clonal expansion is through the interrogation of the diversity of TCR sequences. Antigen-dependent activation results in decreased TCR diversity at the site of insult due to the expansion of T cells with specific TCRs. TCR repertoire diversity is a result of imprecise rearrangement of the TCR alpha and beta loci, resulting in the generation of approximately 100 million unique TCRβ chain sequences in the naïve T cell repertoire of adults^39^. This broad diversity of TCR sequences is significantly reduced following clonal expansion^40^. A significant advantage of our study is the utilization of 5’-end sequencing which allows for the capture of the variable domains of the TCR. We measured the overall TCR diversity of the T cells from the cirrhotic and healthy livers or PBMCs outlined above. As expected, we found that T cells from the peripheral blood had high TCR diversity, whereas T cells from the liver had lower diversity by multiple metrics (Figure 2A). Due to the low numbers of patients in our study, we did not observe a statistically significant difference in TCR diversity between healthy and cirrhotic individuals. To interrogate transcriptional changes in clonally expanded T cells we utilized GLIPH to cluster TCRs that likely respond to the same antigen due to highly similar complementarity-determining region 3 (CDR3) sequences^32^. This algorithm allowed us to identify clonal T cell groups across all samples and by individual disease state (Figure 2B). However, when inspecting the largest clonal groups within any disease status, we noted that there were no clones unique to ND samples. Therefore, we evaluated the transcriptional profile of the of the most expanded T cells clones in ALD, MASH, and PBMC (Figure 2B). We specifically selected the top CD8^+^ T cell clone from each disease for further comparison, as only MASH had a top T cell clone that was both unique and CD4^+^ (Supplemental Figure 3). Although these cells occupied related spaces on the UMAP (Figure 2C), the transcriptional profile of these clonally expanded T cells differed based on disease etiology. The clonally expanded T cells from MASH expressed markers of chronic antigen stimulation (*KLRG1, CD160, CD244, TIGIT, TOX,* and *PDCD1),* whereas ALD and PBMC-derived T cells expressed higher levels of Tcm associated marker *IL7R* (Figure 2D). These results demonstrate that cirrhosis results in the accumulation of activated, clonally expanded T cells in the liver.

**Figure 2:**
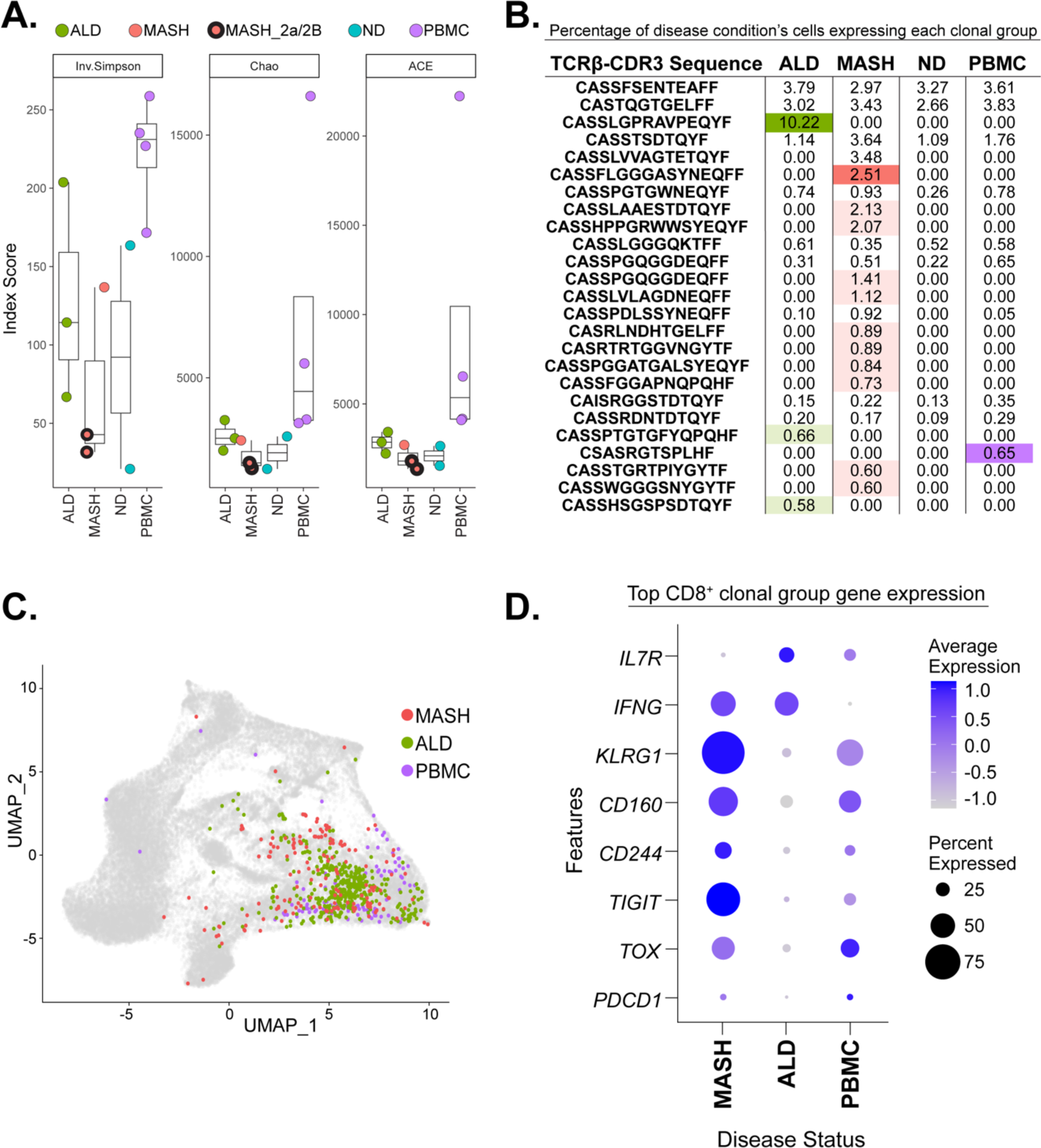
MASH-induced cirrhosis results in the recruitment of clonally expanded, chronically stimulated T cells to the liver. (A) Inverse Simpson (left), Chao (middle), and abundance-based coverage estimator (ACE) (right) indices measuring sample diversity. Boxplots range from 25^th^-75^th^ percentile with a horizonal line at the mean. (B) Top 25 most clonally expanded T cell groups as clustered by GLIPH and ranked by largest cumulative proportion of T cell population from each disease status. Table displays percentage of T cells from disease conditions expressing each clonal group. Darker highlighted percentage indicate most clonally expanded, CD8^+^ T cell clonal group unique to each disease condition. Lighter highlighted percentages indicate other top 25 CD8^+^ clones unique to one disease condition. (C) UMAP plot displaying the most clonally expanded CD8^+^ T cell clonal group unique to each disease condition as identified in B. (D) Gene expression dot plot of selected gene (Features) expression for the most clonally expanded CD8+ T cell clonal group unique to each disease condition. Larger dot size represents larger percent of cells expressing each feature and darker dot color represents the average expression level of cells expressing each feature.

### HFHC-diet feeding induces MASH pathology and accumulation of activated CD8^+^ T cells in the livers of mice

Cirrhosis has unique effects on the immune system both in the liver and systemically^41^, therefore it is unclear if the T cell clonal expansion and/or transcriptional changes in our study were a result of cirrhosis-dependent effects. To answer this question, we examined the profile of liver resident T cells in a well validated mouse model of MASH (high-fat (40%) and high-cholesterol (2%) diet (HFHC)) and evaluated at a timeframe prior to cirrhosis (25-31 weeks). Similar to previous studies ^23,34^, we found that HFHC feeding resulted in increased liver injury and fibrosis (Figure 3A-D). To define the profile of T cells in the liver during HFHC feeding, mice were euthanized, livers were perfused, and liver resident NPC were isolated as previously described^23,34,42^. We found both a significantly increased percentage of total T cells (Figure 3E-G) and increased ratio of CD8^+^ T cells to CD4^+^ T cells in HFHC-fed mice (Figure 3H,I). CXCR6^+^CD8^+^ T cells have been shown to accumulate in the liver of both humans and murine models (western diet and choline deficient HF-diet) of MASH^43^. Consistent with this, we observed a similar increase in the proportion of CXCR6^+^CD44^hi^ CD8^+^ T cells (Figure 4A,B), but not in the CD4^+^ T cell compartment (Figure 4C) of HFHC mice. We also examined the expression of markers associated with chronic stimulation and T cell exhaustion (Tox, Tigit, and Pd1). Individually, the frequency of Tox^+^, Tigit^+^, and Pd1^+^ T cells increased at 27 and 31 weeks on HFHC diet compared to chow controls (Figure 4D). Importantly, HFHC feeding resulted in the emergence of a CD8^+^ T cell population that co-expressed all three markers of chronic TCR stimulation (Figure 4E). Together, these findings demonstrate that HFHC feeding recapitulates the CD8^+^ T cell phenotype in human MASH where the CD8^+^ T cells in the liver express markers associated with activation and exhaustion.

**Figure 3:**
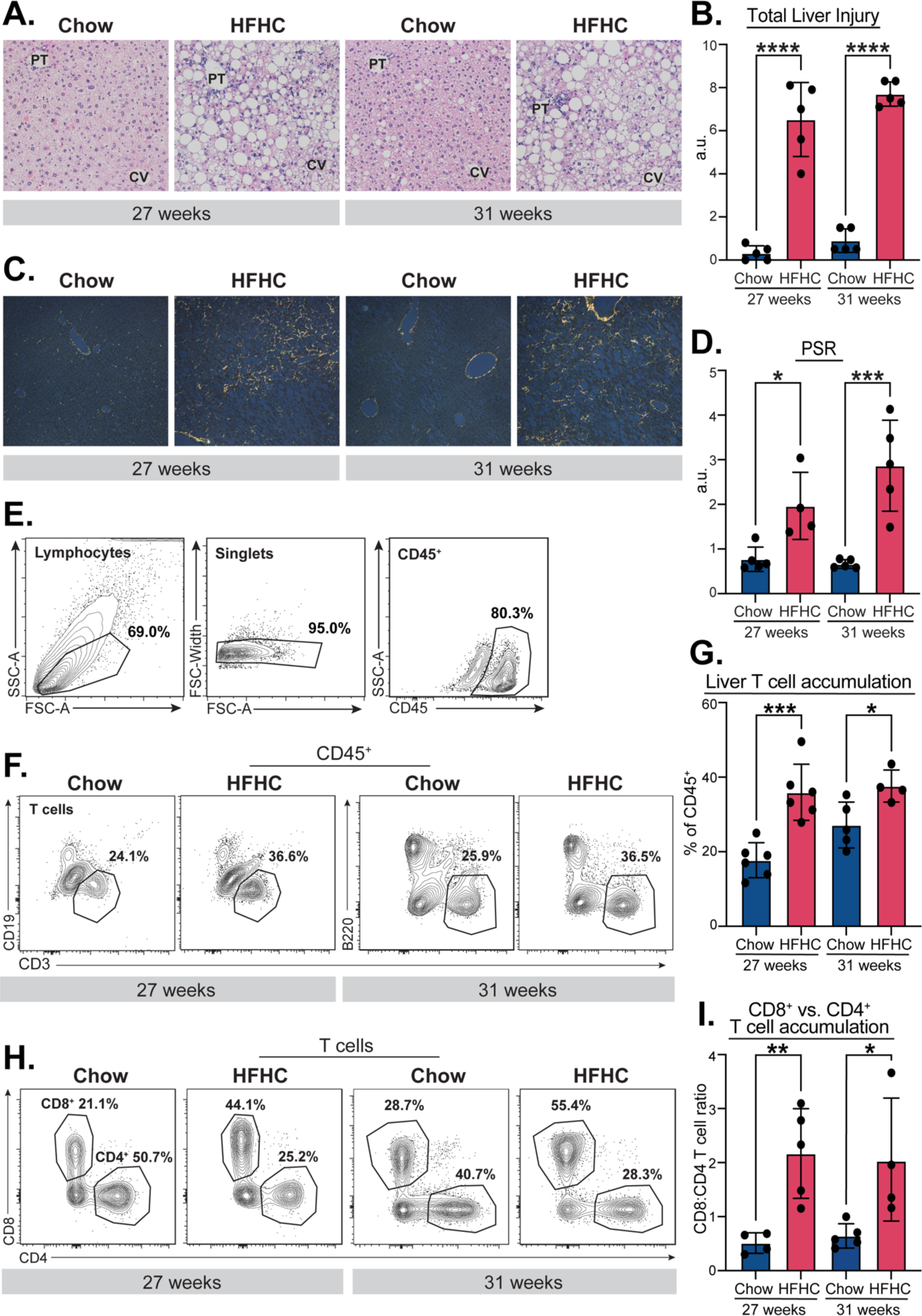
HFHC-feeding recapitulates MASH phenotype in murine models. Mice were fed a chow or HFHC diet for 27 (left) or 31 weeks (right) and investigated for a MASH phenotype. (A) Representative steatosis shown by H&E staining of livers. PT = portal triad, CV = central vein. (B) Total liver injury scoring, as determined by liver cell injury, inflammation, reactive changes, and steatosis from H&E-stained livers. (C) Representative fibrosis shown by PSR staining of livers. (D) Quantification of PSR staining with polarized light. A.U.=arbitrary units. (E) Flow cytometry gating strategy to identify CD45^+^ cells. (F) Representative flow cytometry contour plot showing T cell gating and percentages. (G) Quantification of F. (H) Representative flow cytometry contour plot showing CD8^+^ and CD4^+^ T cell gating and percentages. (I) Quantification of H. n = 4-5/group for quantification above. *P<=0.05, **P<= 0.01, ***P<=0.001.

**Figure 4:**
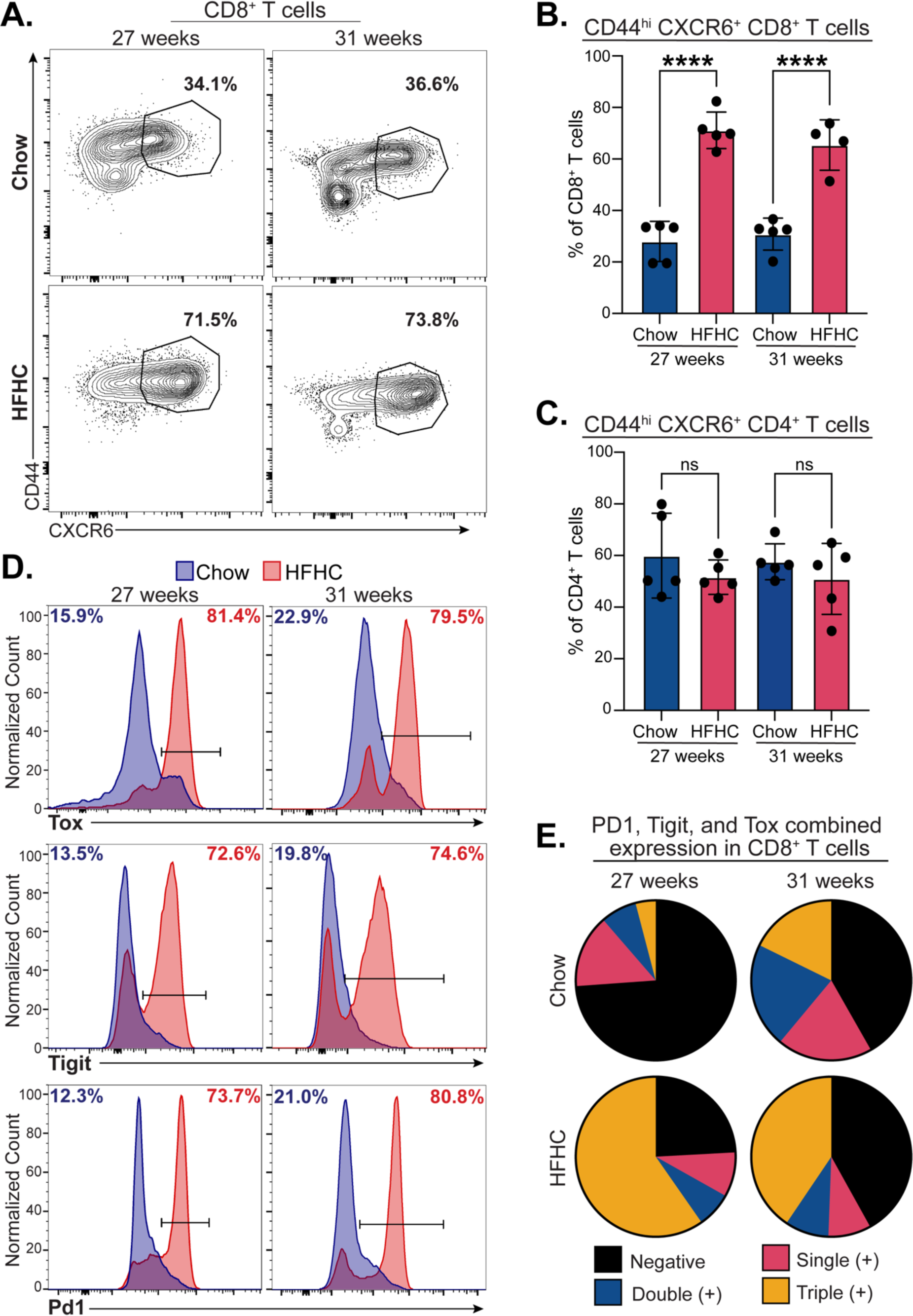
HFHC-feeding leads to increased co-expression of markers associated with chronic stimulation and T cell exhaustion. Mice were fed a chow or HFHC diet for 27 (left) or 31 weeks (right) and investigated via flow cytometry. (A) Representative contour plot showing CD44^hi^CXCR6^+^ of CD8^+^ T cells and percentages. (B) Quantification of A. (C) Quantification of CD44^hi^, CXCR6^+^ CD4^+^ T cells. (D) Representative histograms showing Tox (top), Tigit (middle), or Pd1 (bottom) expression levels and percent positive among CD8^+^ T cells. (E) Pie charts showing the proportion of CD8^+^ T cells that positive for none (Negative), only one (Single (+)), only two (Double (+)), or all three (Triple (+)) of the following markers: Tox, Tigit, and Pd1. n = 4-5/group for quantification and pie charts above. ****P<=0.0001.

### Single cell sequencing reveals novel activated CD8^+^ T cell populations in the liver of MASH

To confirm our flow cytometric findings and identify further differences in T cell phenotype in mice with MASH, liver resident immune cells were enriched for viable T cells and subjected to droplet-based 5’-end scRNA-seq. We profiled T cells from HFHC fed mice and control chow fed mice after 25 and 31 weeks of diet (n = 4/group). Profiled cells were filtered for *Cd3e* expression (Supplemental Figure 4) and then clustered based on transcriptional similarities (Figure 5A,B). Using known markers of T cell functional states, we identified CD8^+^ and CD4^+^ clusters, which were further divided into CD8^+^ Naïve/Tcm, CD8^+^ Effector/Tem, CD8^+^ Exhausted (Tex), NK/NKT, CD4^+^ Naïve/Tcm, CD4^+^ Th1, CD4^+^ Th2, CD4^+^ Th17, and CD4^+^ Treg clusters (Figure 5C,D). Unlike in human cirrhosis, we identified significant changes in the frequency of chow and HFHC derived T cells in each cluster (Figure 5E,F). We found a greater representation of T cells from HFHC mice in CD8^+^ exhausted T cell clusters while other CD8^+^ T cell cluster groups, such as CD8^+^ Naïve/Tcm and CD8^+^ T Effector/Tem, varied in diet origin. In support of our flow cytometry data, the CD4^+^ T cell clusters had a greater representation of chow derived T cells, with the exceptions of Th17 and Treg clusters, which were derived equally from chow and HFHC fed mice and majority from HFHC fed mice, respectively (Figure 5F). Therefore, we show diet-specific differences in the clustering of the profiled T cells and importantly an enrichment of HFHC derived T cells in CD8^+^ Tex clusters.

**Figure 5:**
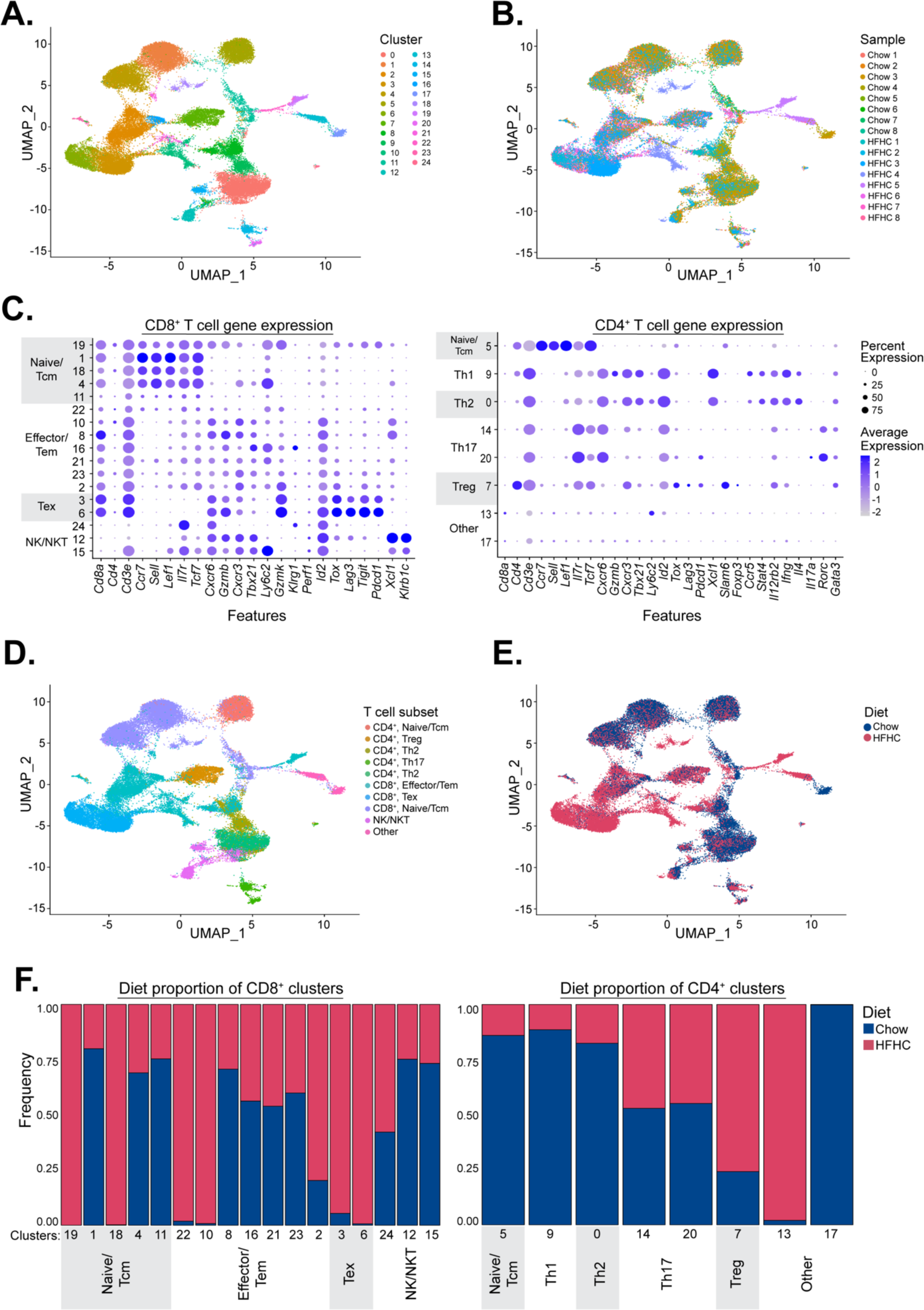
scRNA-sequencing shows T cells derived from HFHC diet fed mouse models of MASH differentially cluster from control mouse T cells. (A) UMAP plot of 49,127 liver resident murine T cells colored by scTriangulate clusters. (B) UMAP plot colored by sample origin of murine T cells after 25 weeks (chow and HFHC sample 5-8) or 31 weeks (chow and HFHC samples 1-4) on diet. (C) Dot plot of selected gene (Features) expression grouped by cluster assignment. Plots are separated based on predominate *Cd8* (left) or *Cd4* (right) expression. Larger dot size represents larger percent of cells expressing each feature and darker dot color represents the average expression level of cells expressing each feature. (D) UMAP plot colored by T cell subset assignment. (E) UMAP plot colored by diet (chow/control or HFHC/MASH) status of T cell sample origin. (F) Bar graph showing proportion of T cells derived from each condition within in clusters for *Cd8*^+^ (left) and *Cd4*^+^ (right).

Next, we examined global gene expression changes between diets, regardless of cluster. Due to the overrepresentation of HFHC derived T cells in CD8^+^ Tex clusters, we investigated the diet-specific expression of exhaustion-related genes, as well as genes associated with naïve and central memory T cells, which are less likely to be associated with T cell exhaustion. Similar to the proteomic profile revealed by flow cytometry, the expression of *Cxcr6* and exhaustion related genes, such as *Pdcd1* (Pd1), *Tigit*, and *Tox,* were all higher in T cells found in the liver of HFHC diet fed mice at both 25 and 31 weeks after starting diet. In contrast, the expression of genes associated with naïve and central memory T cells, such as *Ccr7* and *Sell* (L-selectin/Cd62L), were higher in T cells from livers of chow fed mice (Figure 6A). Since individual transcripts associated with exhaustion were upregulated in HFHC mice, we next sought to inspect inferred pathway differences via Ingenuity Pathway Analysis (IPA). We identified 64 pathways that were significantly upregulated and 31 pathways which that significantly downregulated in T cells from HFHC fed mice (Figure 6B). Pathways enriched in T cells from HFHC mice included ‘TCR signaling’ and ‘Interferon gamma signaling’ while downregulated pathways included ‘IL-15 production’ and ‘Cholesterol Biosynthesis’. Using gene set enrichment analysis (fGSEA) we found, in agreement with IPA, enrichment of gene sets for ‘Interferon Gamma Response’ and ‘TCR Signaling’ in T cells isolated from livers of HFHC fed mice (Figure 6C,D). These findings suggest that, like our findings with CD8^+^ T cells in humans with MASH-induced cirrhosis, the CD8^+^ T cells that accumulate in the liver of mice during the progression of MASH are receiving chronic TCR stimulation.

**Figure 6:**
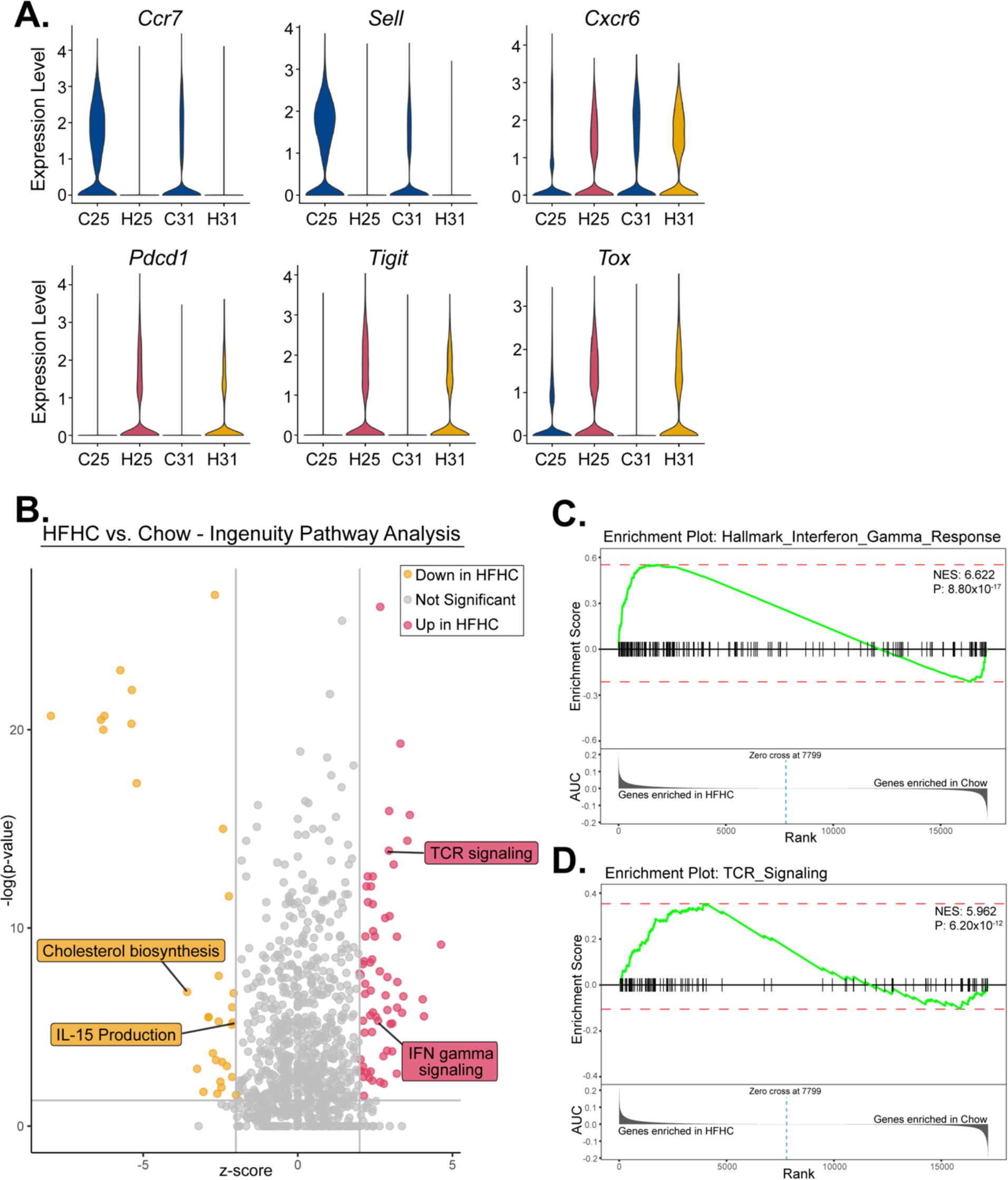
HFHC derived T cells express high levels of transcriptional markers and inferred pathways associated with T cell activation and exhaustion. (A) Violin plots of selected transcriptional marker normalized gene expression: *Ccr7* (Ccr7)*, Sell* (L-Selectin/Cd62L)*, Cxcr6* (Cxcr6)*, Pdcd1* (Pd1)*, Tigit* (Tigit), and *Tox* (Tox), as grouped by diet (C = chow, H = HFHC) and length of diet (25 = 25 weeks, 31 = 31 weeks). (B) Volcano plot of pathways significantly down-regulated (yellow), not significantly different (gray), or significantly up-regulated (red) in HFHC T cells compared to chow T cells as inferred by Ingenuity Pathway Analysis (IPA). (C,D) Gene set enrichment analysis (fGSEA) enrichment plot showing the enrichment score (top) and the area under the curve (AUC) for genes ranked by enrichment in HFHC (left) or chow (right) T cells for (C) the interferon gamma response pathway and (D) the TCR signaling pathway. NES = normalized enrichment score. P = adjusted P-value.

### HFHC feeding causes CD8^+^ T cell clonal expansion in the liver

As the transcriptional profile from liver infiltrating T cells suggests chronic antigenic stimulation, we investigated the clonality of the T cells present in our single-cell dataset using TCR repertoire analysis. First, we examined the overall diversity of the T cells present in HFHC and chow fed mice. In all diversity indices measured, we found a significant decrease in T cell diversity in the T cell populations from the liver of HFHC mice (Figure 7A). We observed a dramatic increase in the expansion of several independent clones from HFHC fed mice at both 25 and 31 weeks of diet (Figure 7B). We next investigated clonally expanded groups containing >20 T cells from HFHC diet-fed mice and noted these groups fell into both the CD8^+^ Tex clusters (67.55%) and the CD8^+^ Effector/Tem clusters (31.81%) (Figure 7C-E). Additionally, as the HFHC clonal groups increased in size, the expression of *Cd160, Cxcr6, Tigit, Tox,* and *Pdcd1* increased and expression of *Il7r* decreased (Figure 7F). Significantly less clonally expanded groups containing >20 T cells were present in the chow mice. The few expanded chow T cells almost exclusively fell into the CD8^+^ Effector/Tem clusters (81.94%) and expressed higher levels of markers related to memory (*Cd69*, *Il7r*) and lower levels of markers related to exhaustion (*Tigit, Tox, Pdcd1*) compared to expanded clones from HFHC mice (Supplemental Figure 5). To expand upon these differences, we investigated the global inferred pathway changes between the clonally expanded T cells found in HFHC mice and those found in chow mice via IPA. From this analysis we identified 91 pathways that were significantly upregulated and 6 pathways that were significantly downregulated in clonally expanded T cells from the livers of HFHC fed mice. T cells from the livers of HFHC mice had increased expression of pathways directly associated with TCR signaling including ‘T cell Exhaustion Signaling’, ‘Interferon gamma signaling’, and ‘nur77 Signaling in T Lymphocytes’ (Figure 8A). Due to the enrichment of the ‘T cell Exhaustion Signaling’ pathway, the notably higher expression of exhaustion-related genes in clonally expanded T cells from the livers of HFHC mice, and the notably higher expression of memory related genes in clonally expanded T cells from the livers of chow mice, we performed fGSEA with a set of genes that are upregulated in exhausted CD8^+^ T cells compared to memory CD8^+^ T cells. This gene set, as well as a gene set of Interferon Gamma Response-related genes, was significantly enriched in the clonally expanded T cells from the livers of HFHC mice (Figure 8B,C). These findings demonstrate that a well validated murine model of MASH results in the accumulation of clonally expanded CD8^+^ T cells in the liver and that the clonally expanded cells express high levels of markers associated with T cell exhaustion. Taken together, our data demonstrate that T cell clonal expansion and thus chronic antigenic stimulation are common events in both human MASH-induced cirrhosis and murine models of MASH. These data provide the framework for further exploration of the causes and consequences of T cell antigen activation in the pathology of MASH.

**Figure 7:**
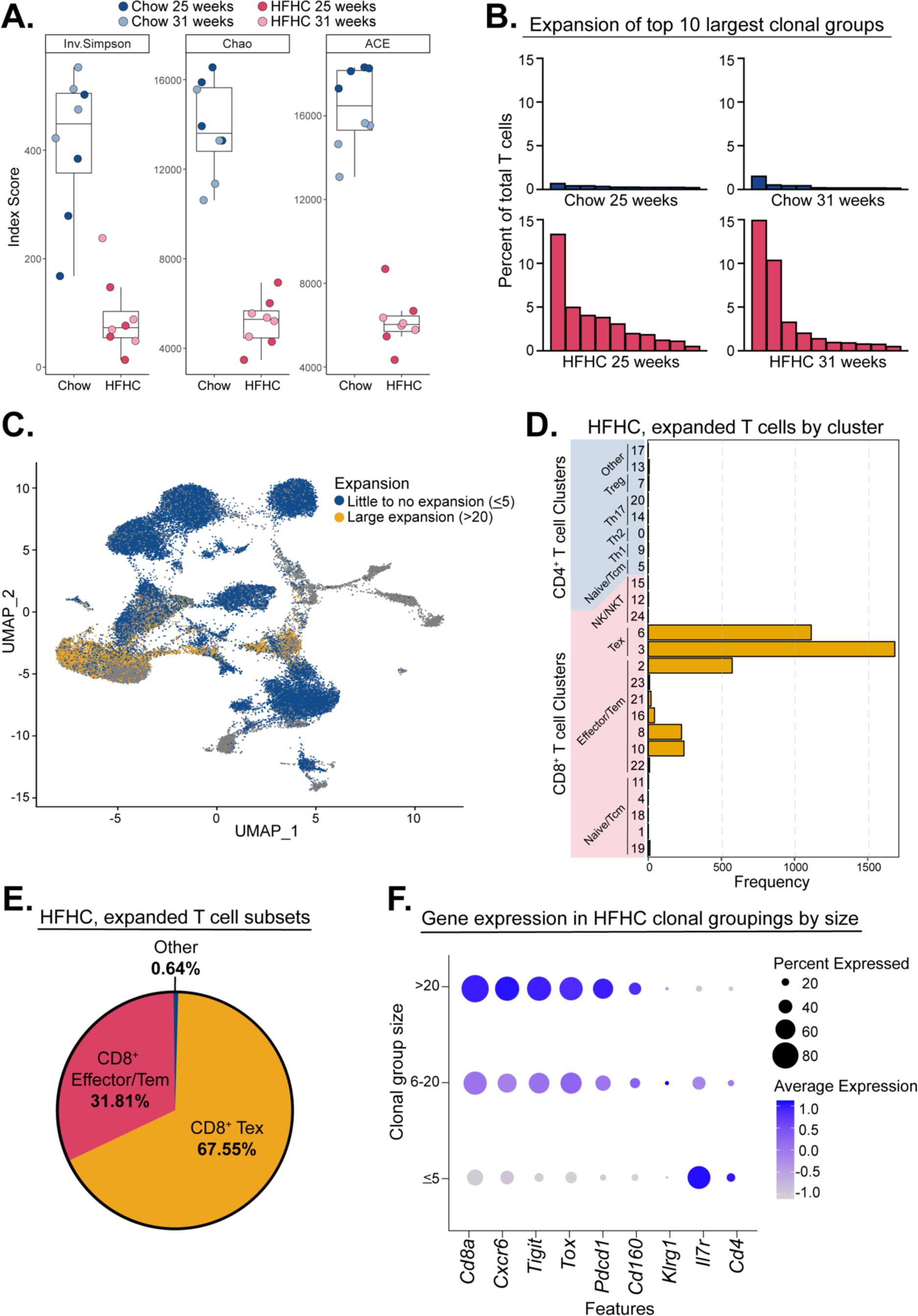
HFHC feeding leads recruitment of clonally expanded CD8^+^ T cell with an exhausted phenotype. (A) Inverse Simpson (left), Chao (middle), and abundance-based coverage estimator (ACE) (right) indices measuring sample diversity. Boxplots range from 25^th^-75^th^ percentile with a horizonal line at the mean. (B) Bar graphs of the percent of total T cells that fall within each of the top 10 most expanded T cell clones from a representative mouse at each diet treatment and timepoint. (C) UMAP plot colored by T cell expansion. Blue = T cells with called TCR sequences shared with a total of 5 or less T cells. Yellow = clonally expanded T cells/T cells with called TCR sequences shared with at least a total of 20 T cells. (D) Bar plot showing the frequency of HFHC, clonally expanded T cells within each cluster assignment. (E) Pie chart showing the proportion of HFHC, clonally expanded T cells that fall into each T cell subset as assigned in Figure 5C,D. (F) Gene expression dot plot of selected gene (Features) expression for all HFHC T cells, grouped by the size of clonal expansion of each T cell. Larger dot size represents larger percent of cells expressing each feature and darker dot color represents the average expression level of cells expressing each feature.

**Figure 8:**
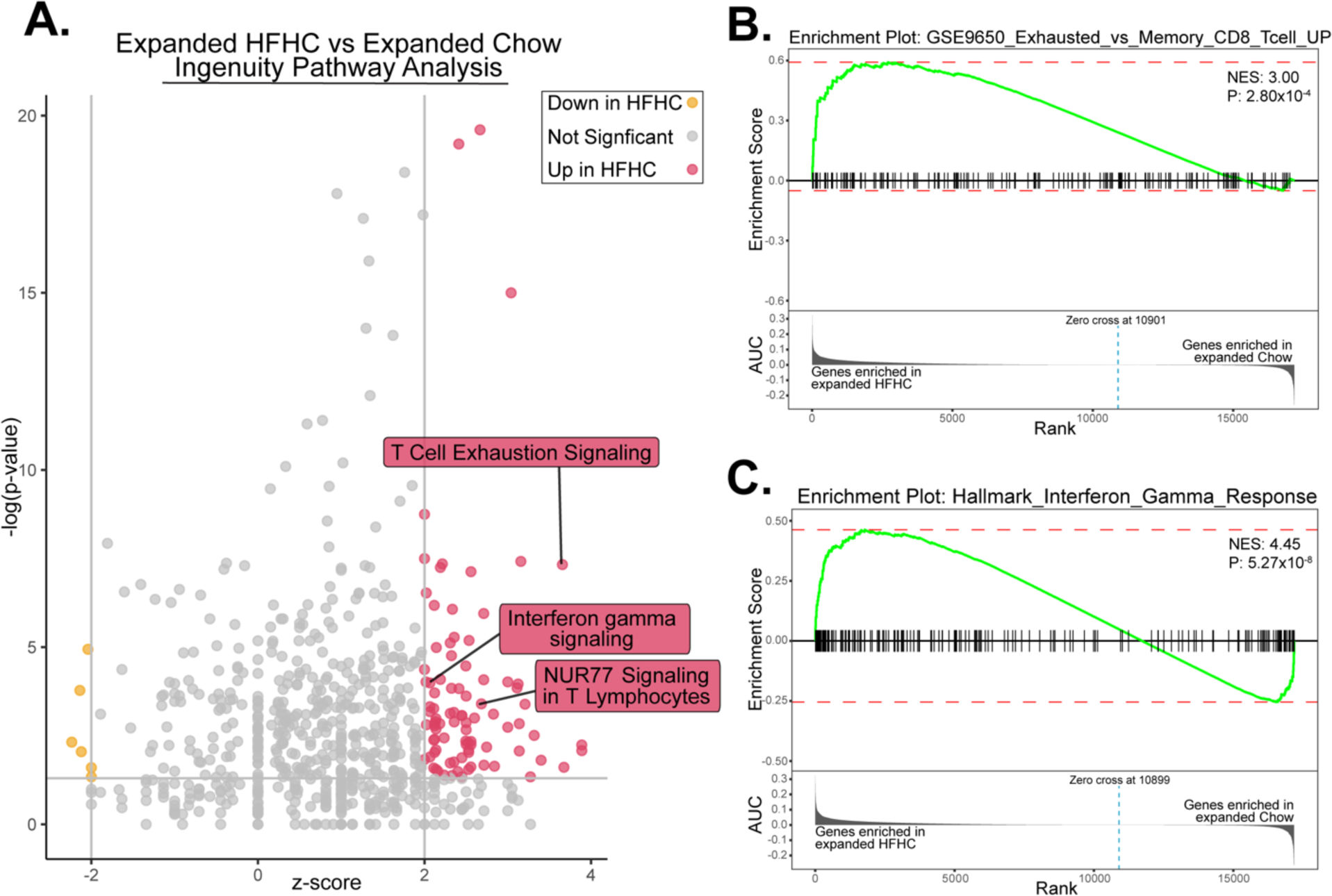
Clonally expanded HFHC derived T cells increase expression of pathways associated with chronic T cell stimulation and exhaustion. (A) Volcano plot of pathways significantly down-regulated (yellow), not significantly different (gray), or significantly up-regulated (red) in clonally expanded HFHC T cells compared to clonally expanded chow T cells as inferred by Ingenuity Pathway Analysis (IPA). (B,C) Gene set enrichment analysis (GSEA) enrichment plot showing the enrichment score (top) and the area under the curve (AUC) for genes ranked by enrichment in clonally expanded HFHC (left) or clonally expanded chow (right) T cells for (B) genes enriched in exhausted CD8^+^ T cells over memory CD8^+^ T cells and (C) the Interferon Response pathway. NES = normalized enrichment score. P = adjusted P-value.

## Discussion

During MASH and other chronic liver diseases, the tolerant nature of the liver is disrupted, and many subsets of activated and pathogenic immune cells are recruited to the liver. This alteration in immune homeostasis is particularly evident during cirrhosis in which immune responses in both the liver and systemically are dysregulated^44^. The removal of these inflammatory immune cells and restoration of normal immune homeostasis are major goals in the treatment of individuals with advanced liver disease^45^. However, the development and application of immune modulators in CLD is hindered by the incomplete understanding of both the functions of immune cells in the progression and resolution of disease and the complex interactions between these cell types in the liver. Thus, deeper characterization of liver infiltrating immune cells in health and disease is required to develop novel therapies for patients with CLD. In our study, we aimed to characterize the T cells that accumulate in the liver at the time of cirrhosis in humans and in a murine model of diet-induced MASH. We demonstrated for the first time, that clonally expanded CD8^+^ T cells accumulate in the livers of both humans and mice with MASH. Thus, we implicate a potential role for antigen activation of CD8^+^ T cells in MASH pathogenesis.

While T cell accumulation has been previously reported by us and others in both humans with MASH-induced cirrhosis and murine MASH models^8,9^, the precise function of T cells in MASH pathogenesis has yet to be resolved. In these studies, we hypothesized that, similar to chronic infection with liver tropic infectious agents such as HCV and HBV, antigen-activated T cells are recruited to the liver to clear an antigenic insult. As previous reports have demonstrated that T cells can be recruited to sites of inflammation in both an antigen-dependent and antigen-independent manner^46–48^, it was plausible that the inflammatory environment in the liver during MASH causes the recruitment and activation of T cells independent of their antigen specificity. However, we did not find this to be the case as we found that the CD8^+^ T cells that accumulated in the liver of humans and mice with MASH expressed high levels of Tigit, Tox, and Pd1 at the protein and transcript levels. Tigit, Tox, and Pd1 are individually and collectively associated with both T cell activation and chronic exhaustion due to stimulation through the TCR^49^. Thus, our results point to T cell accumulation in the liver being a result of antigen driven T cell activation and clonal expansion rather than cytokine driven activation and accumulation.

When presented with peptide in the context of MHC and a proper inflammatory signal, an antigen-specific T cell undergoes clonal expansion, differentiation, and acquisition of effector functions. Thus, a distinguishing feature of antigen-dependent T cell activation is the expansion of T cells with TCRs that recognize the stimulatory antigen. In our study we interrogated the TCR repertoire of liver infiltrating T cells to directly determine if antigen dependent activation occurs in MASH. Through scRNA-seq of liver resident T cells in murine models of MASH, we identified a significant reduction in TCR diversity in the liver infiltrating T cell population due to the presence of large T cell clonal expansions. Thus, these studies confirmed that antigenic stimulation is a driving factor for T cell accumulation in the liver during the progression of murine MASH. These clonally expanded T cells expressed higher levels of *Tox, Tigit,* and *Pdcd1* as clonal size increased and largely clustered with T cells identified as ‘CD8^+^ Tex’. This T cell exhaustion profile outlined in mouse and human MASH is very similar to the profile of T cells that accumulate in the liver in settings where T cell responses are unable to effectively clear their cognate antigen^50^.

A major limitation of our study is that it is currently not possible to identify the stimulatory antigen by exclusively interrogating the TCR sequence. Computational approaches such as TCRMatch^51^ have attempted to predict stimulatory antigens from TCR sequence alone. However, these approaches are limited as they are dependent on a database of previously identified T cell clones. In fact, when we input our TCR sequences into TCRMatch we do not find matching peptides that are biologically relevant. Furthermore, upon inspection of TCR gene usage in murine models of MASH, we observed very little overlap between individual mice (Supplemental Figure 6), suggesting that multiple antigenic targets may cause the T cell activation and accumulation in the liver during MASH. Thus, the antigenic cause(s) of T cell activation in MASH remains unclear and future studies require high-throughput, unbiased approaches^52^ to determine the repertoire of stimulatory antigens in MASH. While elucidating the antigenic causes of clonal expansion in MASH will likely be time consuming, this discovery could clarify the role and function of CD8^+^ T cells in MASH pathogenesis and lead to future targeted MASH treatments, focusing on CD8^+^ T cell function.

## Abbreviations

MASH: metabolic dysfunction-associated steatohepatitis
PD1/*PDCD1*: programmed cell death 1
HCV: hepatitis C virus
ALD: alcohol related liver disease
AIH: autoimmune hepatitis
PBC: primary biliary cholangitis
PSC: primary sclerosing cholangitis
CD: cluster of differentiation
scRNA-seq: single cell mRNA sequencing
NPC: non-parenchymal cells
PBMC: peripheral blood mononuclear cells
TCR: T cell receptor
fGSEA: fast gene set enrichment analysis
IPA: ingenuity pathway analysis
H&E: hematoxylin and eosin
PSR: picrosirius red
ND: non-diseased/no noted liver disease
UMAP: uniform manifold approximation and projection
Tcm: central memory T cell
Trm: resident memory T cell
Tem: effector memory T cell
Th: T helper cell
Treg: T-regulatory cell
Temra: effector memory re-expressing CD45RA T cell
MAIT: mucosal-associated invariant T cell
GLIPH: grouping of lymphocyte interactions by paratope hotspots
CDR3: complementarity-determining region 3
HFHC: high-fat, high-cholesterol
Tex: Exhausted T cell
HBV: hepatitis B virus

## Author Contribution

AECB – study concept and design, data acquisition, analysis and interpretation of data, and drafting of the manuscript. RGS – Analysis and interpretation of the data and critical review of the manuscript. CP – Analysis and interpretation of the data and critical review of the manuscript. DMCD – Acquisition of data and critical review of the manuscript. MEF – Acquisition of data. DJO – Acquisition and analysis of the data. MSK – Sample acquisition and critical revision of the manuscript. BAJT – Data acquisition and critical revision of the manuscript for important intellectual content. AEG – Study concept and design, analysis and interpretation of data, drafting of the manuscript, and statistical analysis. MAB – Study concept and design, acquisition of data, analysis and interpretation of data, obtained funding, drafting of the manuscript, statistical analysis, and study supervision.

