## Supplemental Information for "Antigen-driven CD8^+^ T cell clonal expansion is a prominent feature of MASH in humans and mice"

**Supplementary Table 1: Antibodies used in flow cytometry.**

| TARGET | CLONE | SOURCE |
| --- | --- | --- |
| CD3 | 17A2 | BioLegend |
| CD3e | 145-2C11 | BioLegend |
| CD4 | GK1.5 | BioLegend |
| CD8a | 53-6.7 | BioLegend |
| CD45R (B220) | RA3-6B2 | BioLegend |
| CD19 | 6D5 | BioLegend |
| CD44 | IM7 | BioLegend |
| CD45 | 30-F11 | BioLegend |
| CD186 (CXCR6) | SA051D1 | BioLegend |
| Tox | TXRX10 | eBioscience |
| Tigit | 1G9 | BioLegend |
| CD279 (PD1) | 29F.1A12 | BioLegend |

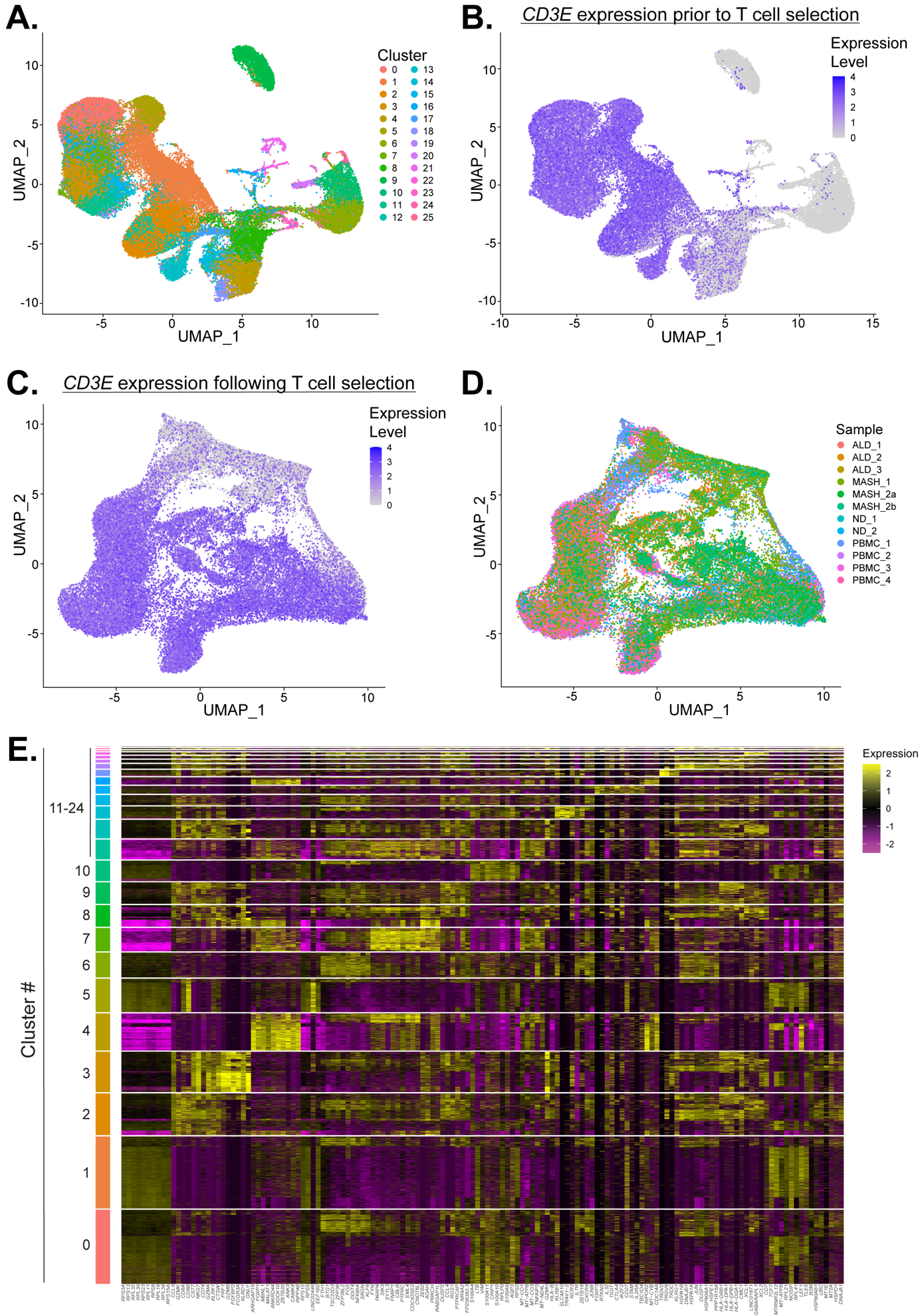

**Supplemental Figure 1: scRNA-sequencing reveals transcriptionally distinct T cell clusters.**

(A) UMAP plot of 61,318 cells colored by scTriangulate clusters prior filtering on *CD3E* expression. (B) Feature plot representation of *CD3E* expression levels across UMAP plot prior to filtering on *CD3E* expression. (C) Feature plot representation of *CD3E* expression levels across UMAP plot of 51,624 cells following exclusion of clusters devoid of *CD3E*. (D) UMAP plot colored by sample ID. (E) Heatmap of top 10 differentially expressed genes by cluster. Yellow represents higher transcript expression while purple represents lower transcript expression.

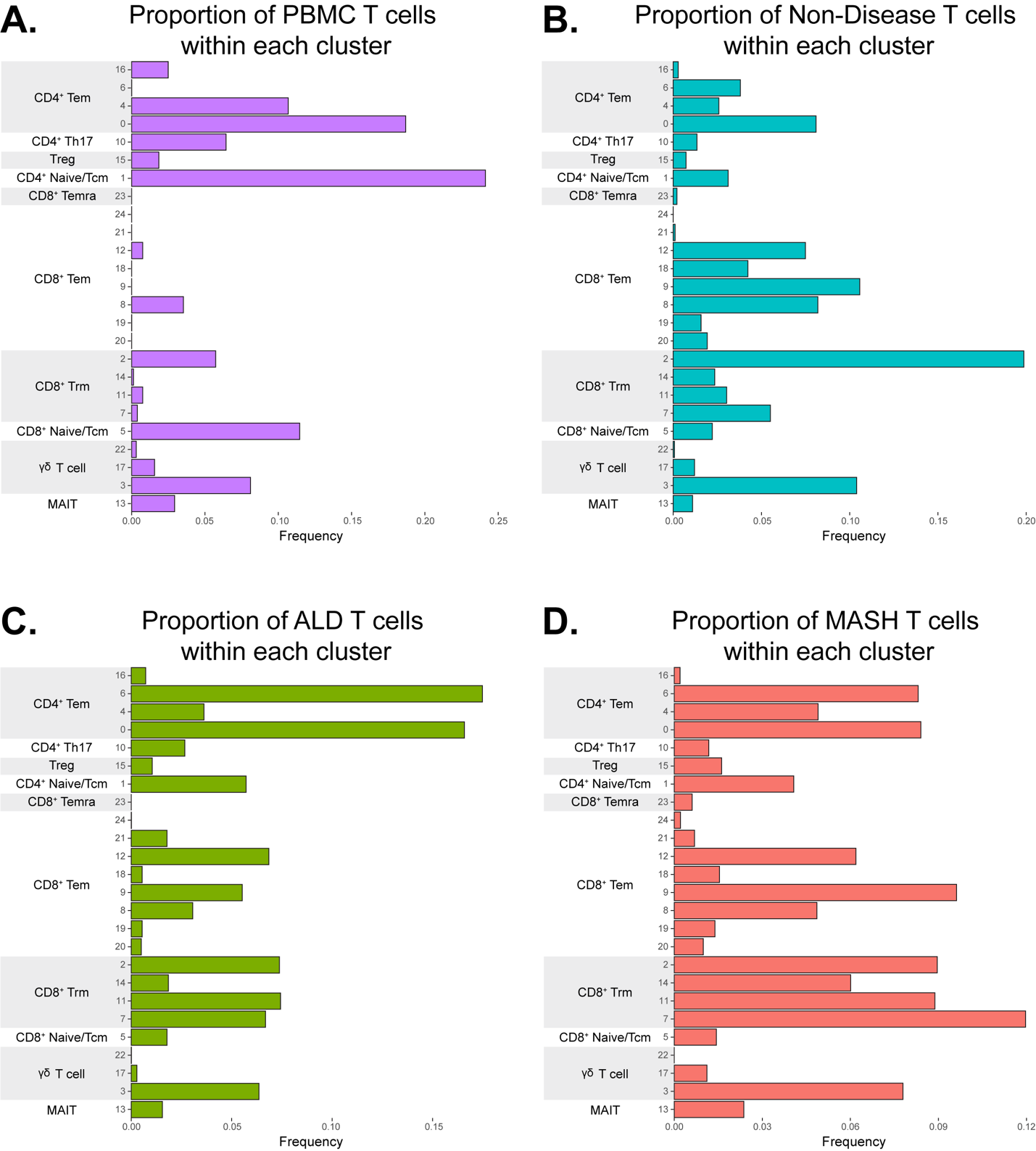

**Supplemental Figure 2: Proportion of T cells within each cluster by disease status.**

Proportion of T cells within each cluster of T cells derived from (A) PBMC samples, (B) Non-diseased (ND) samples, (C) ALD samples, and (D) MASH samples.

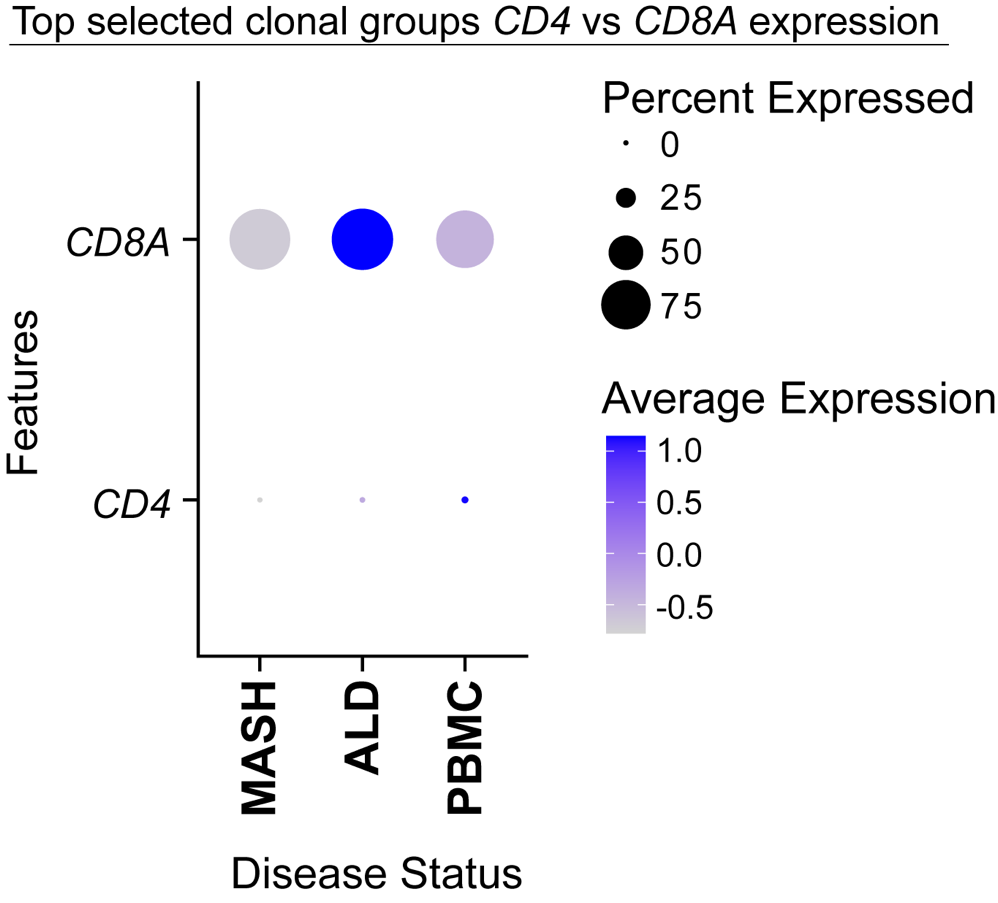

**Supplemental Figure 3: *CD4* vs *CD8A* expression of the top selected T cell clonal groups.**

Gene expression dot plot of *CD4* and *CD8A* expression for the top T cell clonal group that is unique and CD8^+^ for each disease condition. Larger dot size represents larger percent of cells expressing each feature and darker dot color represents the average expression level of cells expressing each feature.

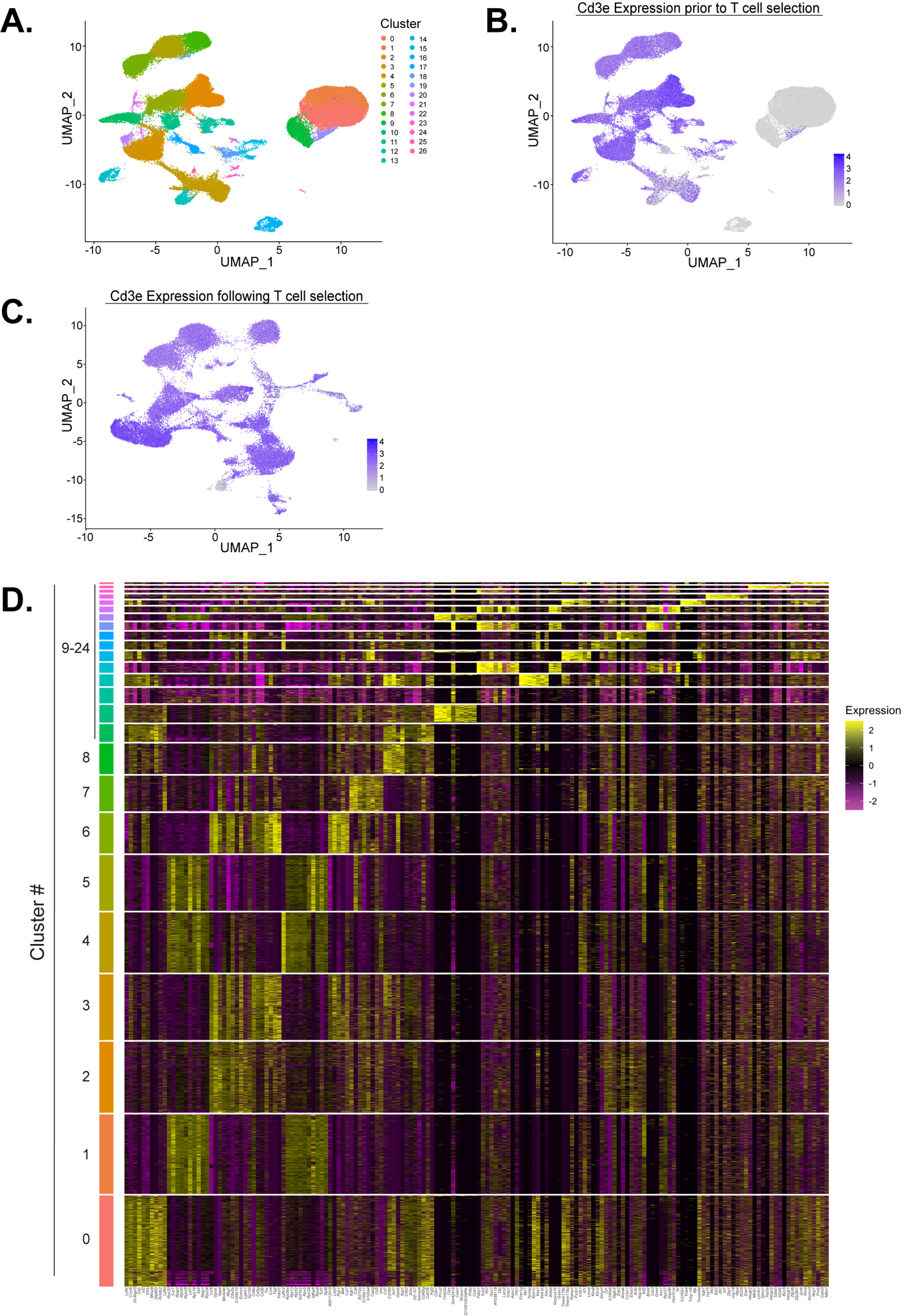
**Supplemental Figure 4: scRNA-sequencing reveals transcriptionally distinct T cell clusters following *Cd3e* expression enrichment of murine T cells.**

(A) UMAP plot of 108,825 cells colored by scTriangulate clusters prior filtering on *Cd3e* expression. (B) Feature plot of *Cd3e* expression levels across UMAP plot prior to filtering on *Cd3e* expression. (C) Feature plot of *Cd3e* expression levels across UMAP plot of 49,127 cells following exclusion of clusters devoid of *Cd3e* expression. (D) Heatmap of top 10 differentially expressed genes by cluster. Yellow represents higher transcript expression while purple represents lower transcript expression.

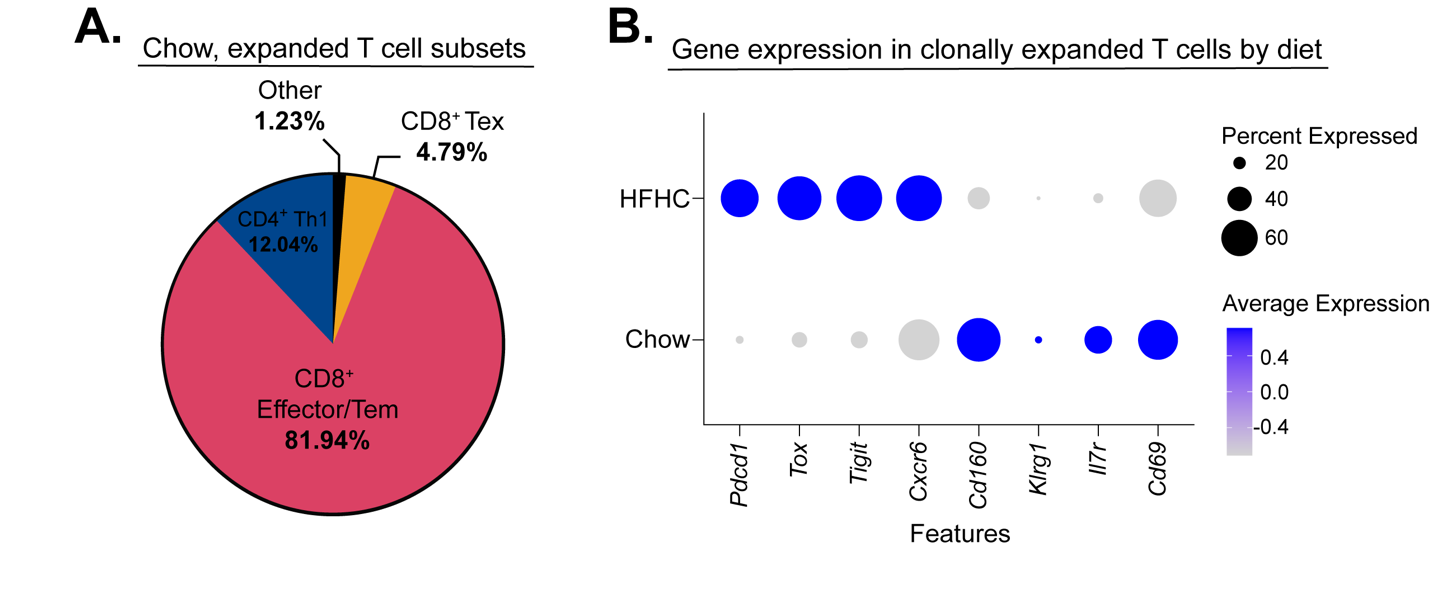

**Supplemental Figure 5: Clonally expanded T cells from chow fed mice do not have an exhausted phenotype.**

(A) Pie chart showing the proportion of chow, clonally expanded T cells that fall into each T cell subset as assigned in Figure 5C,D. (B) Gene expression dot plot of selected gene (Features) expression for all clonally expanded T cells (>20 in clonal group), grouped diet origin. Larger dot size represents larger percent of cells expressing each feature and darker dot color represents the average expression level of cells expressing each feature.

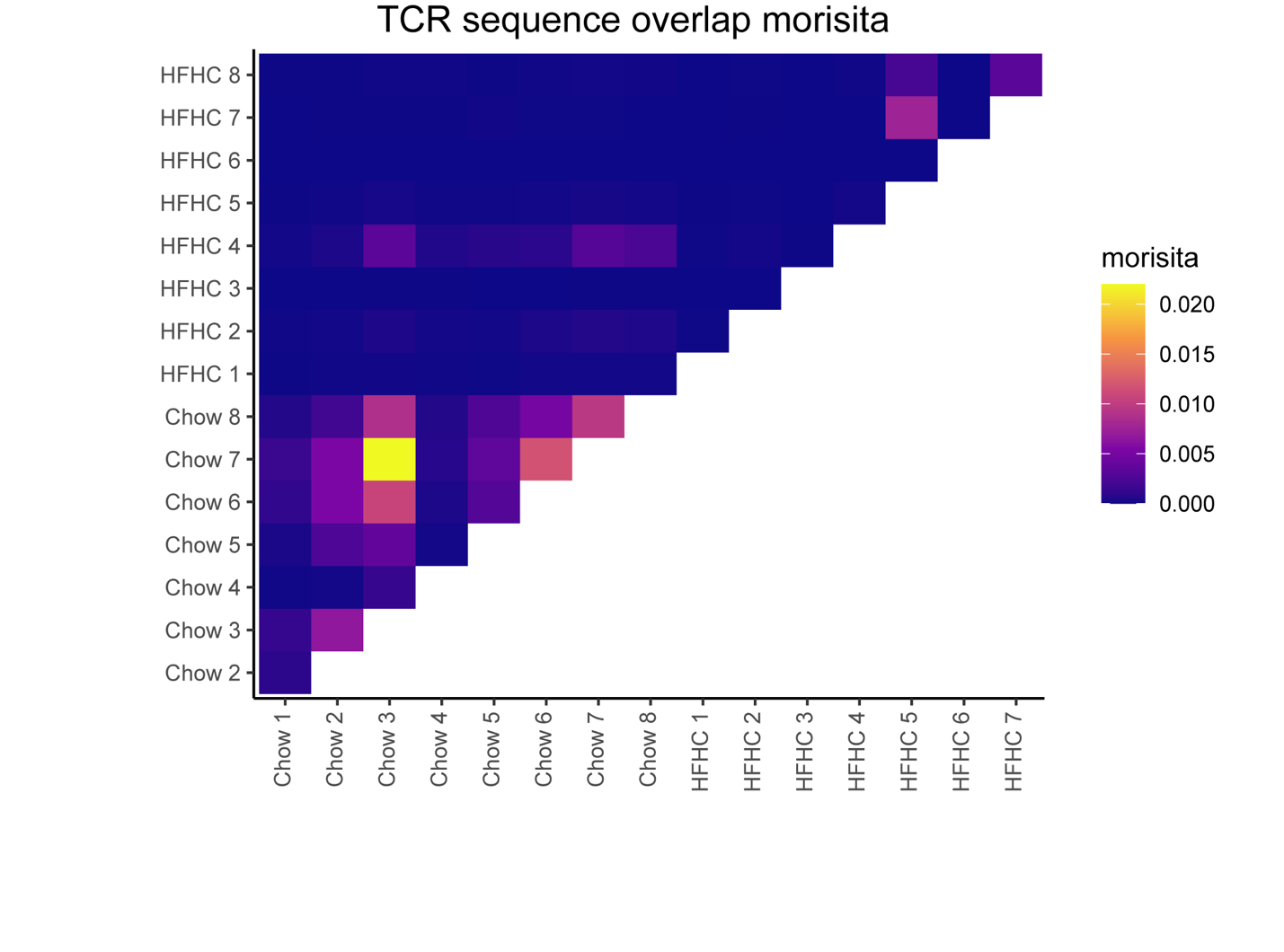

**Supplemental Figure 6: Minimal TCR sequence overlap exists between individual HFHC mice.**

Overlap of TCR sequence usage between samples as calculated by the morisita method on scRepertoire. Yellow = higher overlap between two samples, purple = lower overlap between two samples.
